# Diversity and abundance of pestiferous fruit flies (Diptera: Tephritidae) infesting cucurbit crops in Morogoro, eastern-central Tanzania

**DOI:** 10.1101/2023.04.20.537624

**Authors:** Petronila Tarimo, Sija Kabota, Maulid Mwatawala, Ramadhan Majubwa, Abdul Kudra, Massimiliano Virgilio, Marc De Meyer

## Abstract

Fruit flies represent a significant threat to cucurbit production. They inflict about 30% to 100% losses on cucurbit crops. The aim of this study was to characterize the community of fruit flies infesting cucurbit crops in Morogoro, Tanzania. We investigate the diversity and abundance of fruit fly species infesting cucumber, watermelon, and squash across the two agroecological zones of the Morogoro using three attractants (Cue Lure, Zingerone and Biolure). The attractants were deployed for 16 weeks from June to November 2020.

In total, 12017 fruit fly specimens were collected. Of these, 77.23% were collected from the mountainous and the remaining 22.77% from the plateau zone. All specimens belonged to the genera *Dacus, Zeugodacus* and *Bactrocera*. *Zeugodacus cucurbitae* was the most abundant species (77.6%) while the remaining species constituted 22.4%. The mountainous zone harboured a significantly higher number of fruit fly species than the plateau zone. The abundance of fruit flies was significantly influenced by altitudes and seasons. The study characterized the community of fruit flies in Morogoro and confirms the prevalence of Z. *cucurbitae, D. bivittatus and D. punctatifrons* as prominent species in this region. Therefore, management programs should focus on containing these species regardless of the agroecological zones and seasons.

## 1. Introduction

Fruit flies (Diptera: Tephritidae] are recognized as important pests of fruit and vegetables, causing high economic losses to farmers worldwide [13,17, 50, 66]. According to [19] and [45], fruit flies account for about 30-100% of fruit and vegetable losses in sub-Saharan Africa (SSA) [19, 66]. The SSA alone harbours more than 1,000 species of fruit flies, of which 29 are considered of major economic significance to fruit and vegetable crops [53, 62]. The vast majority of these are indigenous to SSA while a few are of exotic origin [2, 35, 46].

The impacts of fruit fly infestation on horticultural crops in SSA continue to be a major challenge to farmers [2, 35, 46], and the biodiversity monitoring studies have been few and have contributed to the further development of management strategies [2, 29, 30, 42, 48, 54]. In general, there is a fair amount of fruit fly diversity studies in SSA [23, 24, 43, 48, 62], but limited ones on cucurbit crops specifically. Nevertheless, most of these studies on cucurbits were restricted to a narrow range of ecological zones, attractants, limited species richness estimation procedures and a few species, and mostly used para-pheromones and food baits or reared fruits [30, 43, 48, 62].

So far only a few studies by [43] determined the biodiversity of fruit flies using combined methods and procedures of biodiversity assessments. Understanding biodiversity changes across ecological zones requires robust observations and reliable biodiversity assessments based on sampling completeness and combined estimation procedures of species richness [5, 12, 16]. Such biodiversity inventory designed around sampling completeness and robust estimation procedures often provides accurate characterization of fruit fly communities across ecological zones [16, 43].

Moreover, satisfactory efforts must be reached to realize the sampling completeness. According to [6], sampling completeness can be determined using extrapolating species accumulative curves (SACs) and non-parametric estimators of species richness. SACs allow quantification of both within-inventory efficacy, sampling completeness and estimation of the minimum sampling efforts required to obtain a sample that reflects a true community [asymptotic species richness) [10, 34].

Based on SACs, sampling effort is expressed as the number of pooled individuals necessary to reach the asymptotic species richness [8,43]. On the other hand, based on non-parametric estimators of species richness, sampling effort is expressed as the ratio between the observed and theoretical species richness [7, 39, 43]. Species diversity uses mathematical expression indices to represent a community’s ecological diversity based on species richness and evenness [39].

In this study, we determined the diversity and richness of fruit fly species infesting cucurbit crops using diversity indices and non-parametric estimates of species richness in order to develop ad hoc and bespoken management programs for fruit fly species infesting cucurbit crops in Morogoro, Tanzania.

## 2. Materials and Methods

### 2.1. Study site

Studies on the diversity and abundance of pestiferous fruit flies attacking cucurbit crops were conducted from March to November 2020 in Morogoro Region, eastern-central Tanzania. The Morogoro Region is in the transition zone between the bimodal and unimodal rainfall belts at S5°58’-S10°0’ and E35°25’-E38°30’ [60]. Study sites (plateau and mountainous) were selected based on their differences in climatic conditions and farming practices within the Morogoro region (Table 1).

**Table 1.** Agroecological zones of Morogoro Region.

| Agroecological zone | Characteristics |
| --- | --- |
| **Mountainous zone** | Altitude: > 600 meters asl; Average rainfall: 800 mm – 2500 mm p.a, and an annual average temperature of 29°C.<br>The landscape is mainly under agroecological farming |
| **Plateau** | Altitude: 300-600 meters asl; Average rainfall: 700 mm – 1200 mm p.a, the yearly average temperature of ~24°C<br>The landscape is mainly under conventional farming |

Source: [60]

Five experimental plots located at least 1 km apart were established in each agroecological zone (Table 2). Three cucurbitaceous hosts were established per plot, viz; cucumber (*Cucumis sativus* L.), variety “Ashley,” watermelon (*Citrullus lanatus* (Thunb.) Matsum. & Nakai), variety “Sugar baby,” and squash (*Cucurbita moschata* D.) variety “Waltham”. Cucumber, watermelon, and squash were each planted on an area of 1 120 m^2^ (70 m x 16 m) plot size at a spacing of 50 cm x 60 cm, 1 m × 1.5 m and 1 m × 1.5 m respectively. The study was conducted for two cucurbit cropping seasons (from the end of June to the end of August 2020 as cropping season I and from the end of September to early November 2020 as cropping season II).

**Table 2.**
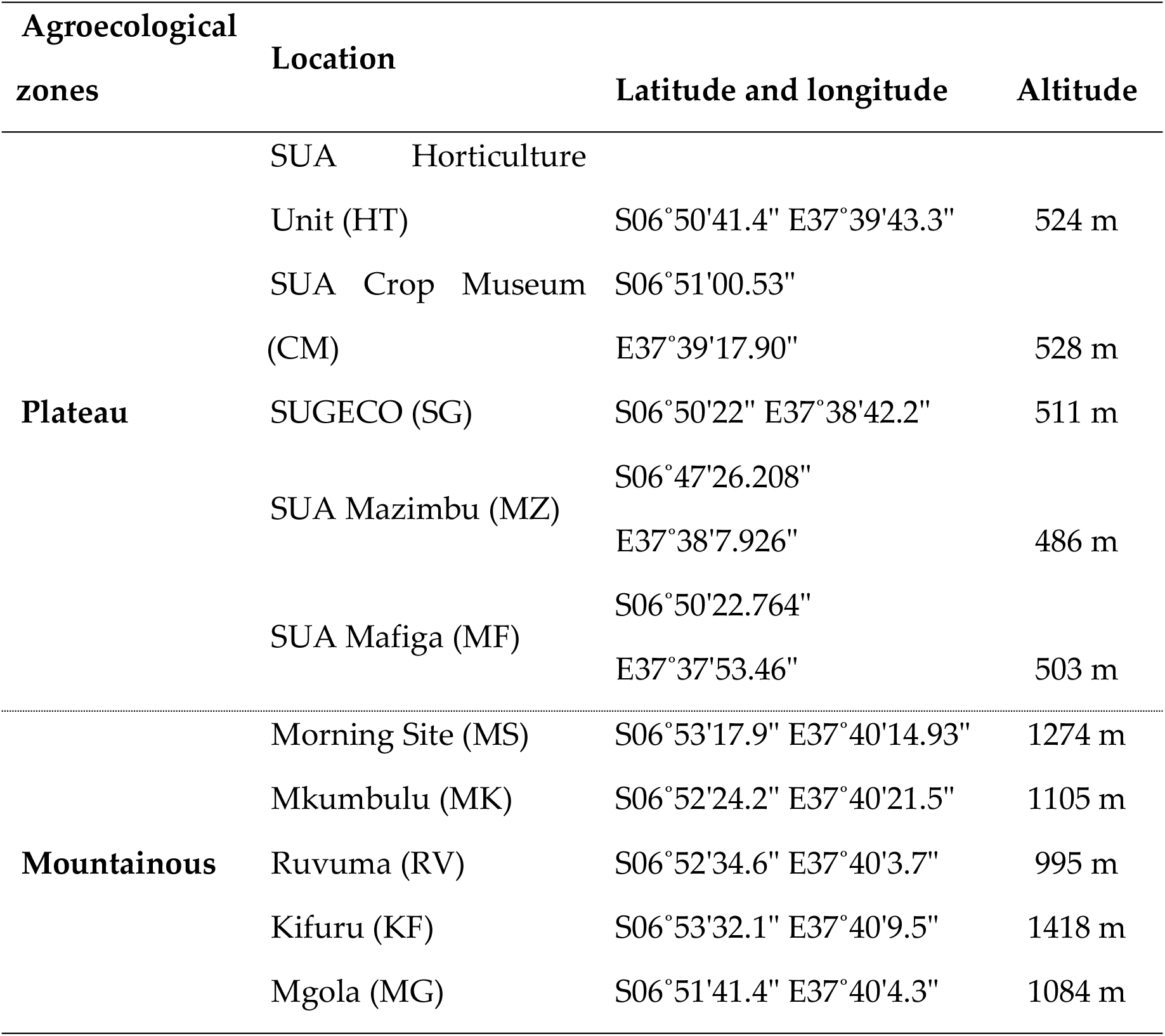
Experimental field locations in two agroecological zones of Morogoro.

### 2.2. Attractants and trapping

Fruit flies were trapped in the 10 established cucurbit plots using clusters of three Tephri-Traps (Sorygar, Spain) each baited with commercially available Cue-lure (CL)(Scentry Biologicals, Billings, Montana, USA), Zingerone (ZN)(Sigma-Aldrich, St. Louis, Missouri, USA) and Bio-Lure (BL)(Chempac (Pty) Ltd, Suterra LLC, USA) for a duration of eight weeks in each season. To minimise lure interference, within each plot, traps were placed at distances of at least 20 m. The lures were selected to attract taxa from the genera *Dacus* and *Zeugodacus*. CL and ZN are male-specific while BL is a general attractant mainly for female fruit flies.

The zingerone was available in crystalline form, which was melted at 40°C in a glass beaker using a heated bath method [26,37]. Once in liquid form, ZN was applied with a graduated pipette to individual 1 x 3 cm cotton dental wicks [52]. CL was available as a plug while BL was supplied in a sachet. Traps were set following the guidelines provided by IAEA [27]. A set of three traps, each baited with one of the attractants, was established in each experimental plot. Each trap was baited with one of the attractants placed on a tree branch or held on a wooden pole at a height of 1.5 m above the ground.

An insecticide dichlorvos (DDVP) strip (Vaportape II; Hercon Environmental, Emingsville, PA, USA) was placed at the bottom of each trap to kill trapped insects. Sticky glue (Tanglefoot) was applied at the base of branches or poles to prevent ants from accessing traps. Traps were inspected once a week, and specimens from each trap were placed in a vial to represent one sample unit. Each vial was marked with unique numbers to represent one sample corresponding to a data sheet with details on date, location, and lure. In order to minimize location bias traps were randomly replaced within the plot after each inspection. Attractants and insecticide strips were replaced every four weeks. Collected specimens were transported to the entomology laboratory at the Sokoine University of Agriculture and preserved in 70% ethanol prior to sorting and identification.

### 2.3. Data collection and identification

Trap catches were recorded as the number of fruit flies per trap per week (FTW). Fruit fly identification was conducted by Petronila P. Tarimo at the Sokoine University of Agriculture entomology laboratory with the aid of a binocular stereomicroscope using keys and characters presented by [3, 63, 64, 65]. Some specimens were sent to the Royal Museum for Central Africa (RMCA] for further identification and confirmation by Marc De Meyer.

### 2.4. Statistical analysis

In this study, sampling efforts in each agroecological zone were evaluated using three non-parametric abundance-based Estimators (ABE) of species richness (Abundance-based Coverage Estimator (ACE), Chao 1, and Jackknife 1) following the protocol described by [7], and Integrated curves (rarefaction (interpolation) and prediction (extrapolation)) as described by [8]. Moreover, the determination of species diversity between the two agroecological zones based on the number of species and relative abundance was done using both α and β diversity indices. Only catches from traps baited with bio-lure were used. This is because the bio-lure is a neutral lure that does not discriminate between fly species and hence allowed the actual comparison of fruit fly species diversity based on the relative abundance of each species.

The α-diversities between the two agroecological zones was assessed using the index of Shannon 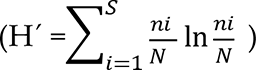 [55], Inverse Simpsons 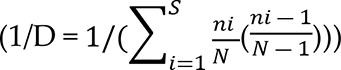[57], and species richness index of Margalef [inine] [38] as well as the evenness index of Pielou 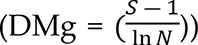[36]. The Shannon and Inverse Simpsons indices take into account the number of species and the relative abundance of each species in a community [43,44].

They both take into account all weight extremes of species abundances ranging from dominance to rareness and therefore, their diversity values are not greatly affected by sample size but are particularly dependent on the extent to which each species is represented among the samples [20]. The species richness by Margalef index was used to highlight the most species-rich zone. The computations of indices were done using the vegan package in R-statistical software.

Data on fruit fly richness, evenness, Inverse Simpson and Shannon were first checked for normality using the Shapiro–Wilk test and for homogeneity using Cochran’s test in the ‘GAD’ package in R. A two-way Analysis of variance (ANOVA) was performed following the general ANOVA Design (GAD) as described by [59] to determine the effects of season and agroecological zones on abundances, richness, evenness, Inverse Simpson, and Shannon values of dominant fruit fly species. Post -hoc comparisons were also performed on the selected model based on the C.test results using the Student-Newman-Keuls (SNK) at alpha =0.05.

The β-diversity was determined using the Sorensen and Jaccard indices. The two indices are based solely on species incidence (presence-absence) data, i.e. the number of species shared by two communities and the number of species unique to each [9,10, 12, 25]. The Sørensen index gives double weight to the shared species between the two communities i.e it weights matches in species composition between the two communities more heavily than mismatches while the Jaccard gives more weight to mismatches in species composition between the two communities [9,12]. The indices are equal to 1 when there is a complete similarity between communities and approach 0 when the communities have no species in common. All statistical models and computations were performed using R -statistical software version 4.1.1 [58]

## 3. Results

### 3.1. Sampling efforts

Expressed as a percentage of the ratio between the observed species richness to the average abundance-based showed that the sampling effort was sufficient to catch 85.56% and 92.67% of all fruit fly species infesting cucurbit crops in the mountainous and plateau zones respectively (Table 3). Moreover, based on individual-based rarefaction and extrapolated curves the asymptote of species richness was reached in all agroecological zones at the pooled value of 9700 and 3200 individuals for the mountainous and plateau respectively (Fig 1). These curves further indicated that the diversity of fruit fly species collected from each of the agroecological zones reflects a more than 85% true fruit fly community of the Morogoro region.

**Fig 1.**
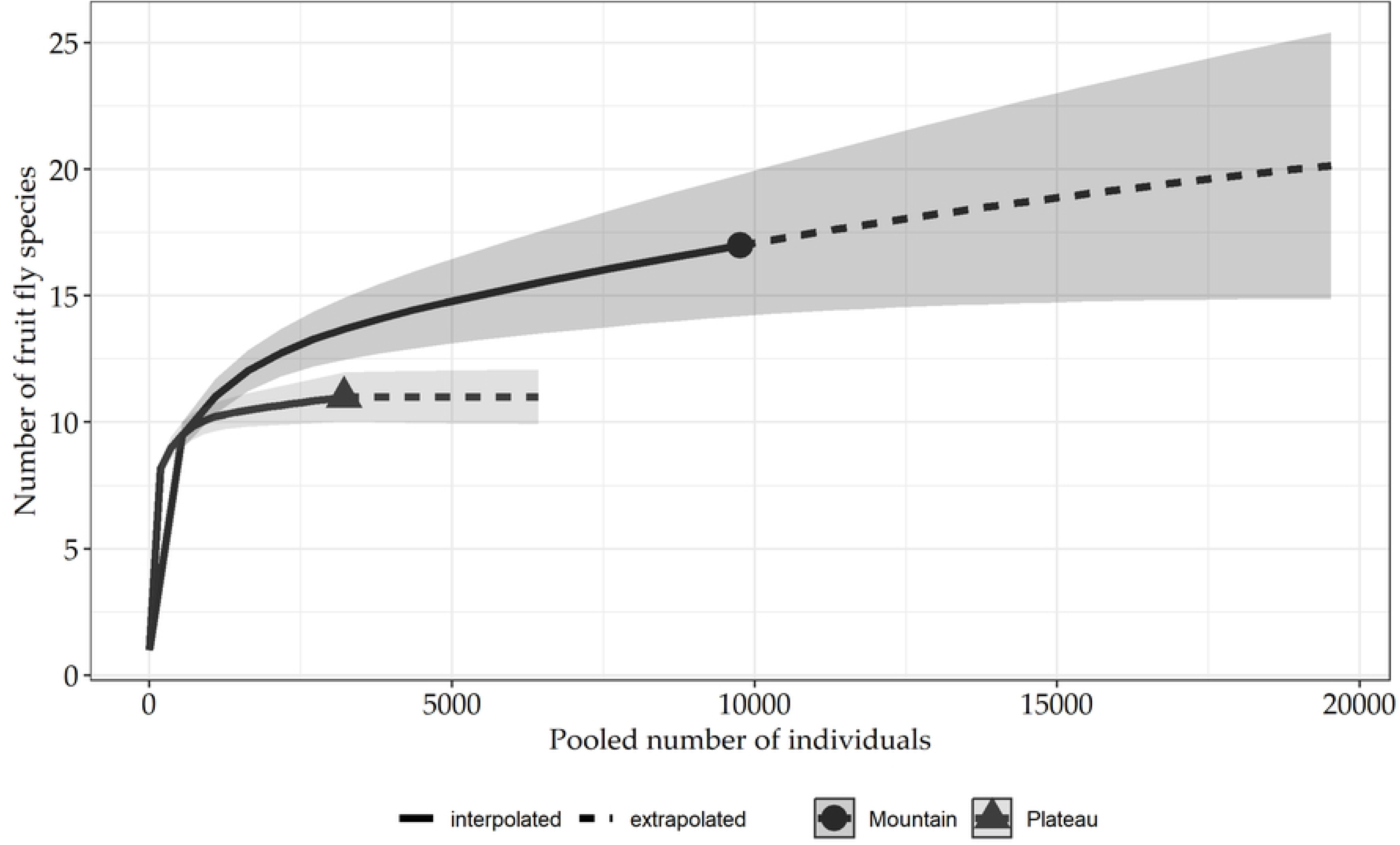
Rarefaction and extrapolation curves based on the number of fruit fly individuals collected in two of the agroecological zones of Morogoro, from April to November 2020.

**Table 3.** Estimators of species richness based on the abundance of fruit flies

| Richness Estimators | Mountainous zone | Plateau zone |
| --- | --- | --- |
| Sobs | 17 | 11 |
| ACE means | 18.28 | 11.2 |
| Chao1 | 20.33 | 11.8 |
| Jack 1 | 21 | 12.6 |
| Mean of the three ABE | 19.87 | 11.87 |
| Sampling efforts (%) | 85.56 | 92.67 |

KEY: Sobs: number of species observed generated per Mao Tau.
ABE: Abundance-Based Estimators.
ACE: Abundance-based Coverage Estimator.

### 3.2. Fruit flies’ abundance

In total, **12,017** fruit fly specimens were collected from the ten cucurbit plots across the two agroecological zones (S 1 Table). Specimens belonged to three genera of *Dacus, Zeugodacus* and *Bactrocera*. Of the total specimens collected; 77.23% were from the mountainous zone and the remaining 22.77% were from the plateau zone. Species from the genus *Zeugodacus* were the most abundant and accounted for 77.6% of the total specimens, while species from the *Dacus* and *Bactrocera* genera constituted the remaining 22.4% (S 1 Table; Fig 2).

**Fig 2.**
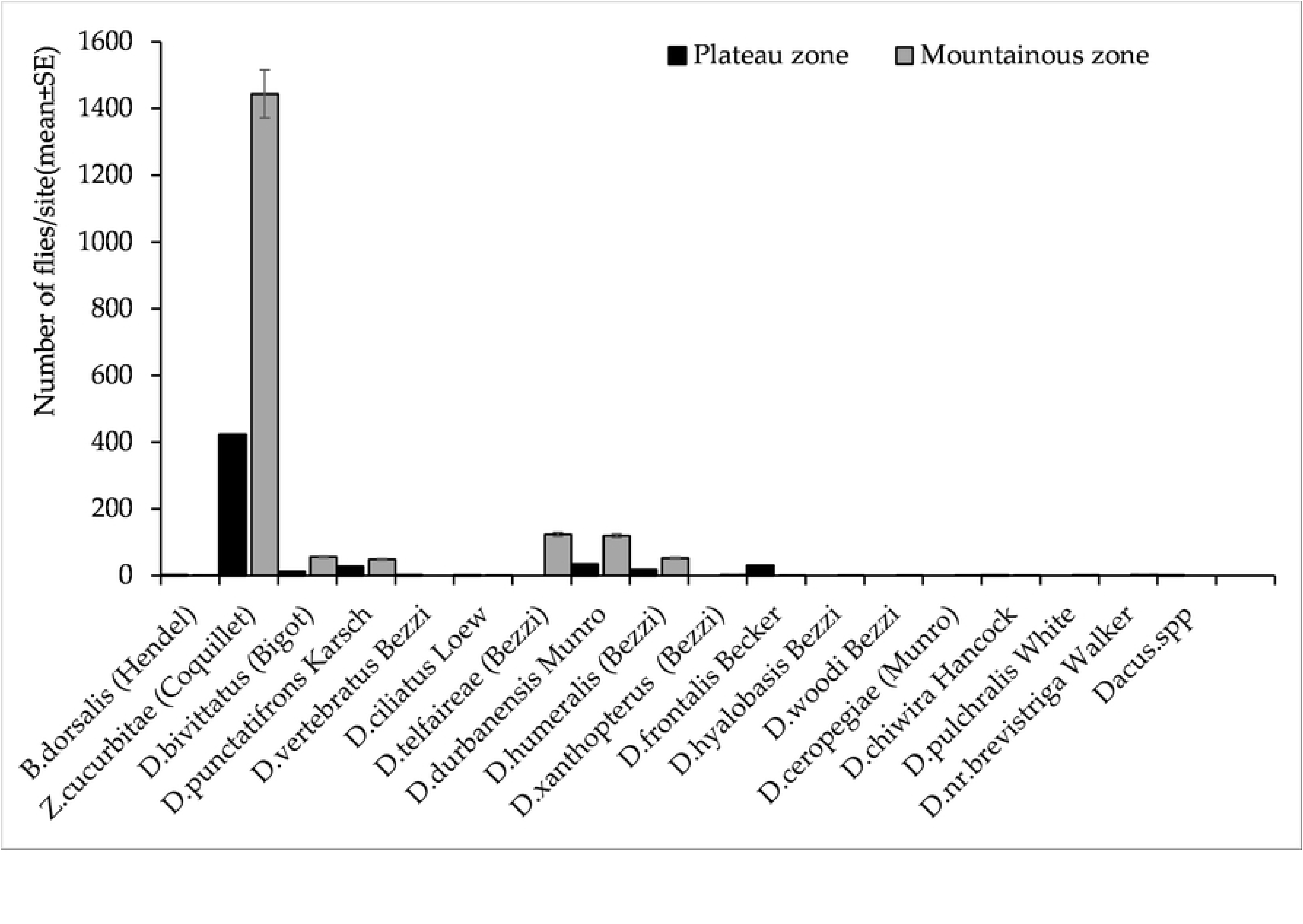
The abundance of fruit fly species infesting cucurbit crops in the mountainous zone of Morogoro, Tanzania from March to October 2020.

Of the 12,017 fruit fly specimens collected, *Z. cucurbitae* was the most abundant fly, followed by *D. durbanensis* Munro and *D. telfaireae* (Bezzi) (S 2 Table). Results also show that *D. durbanensis, D. humeralis* (Bezzi) and *D. frontalis* Becker were more abundant in zingerone compared with Biolure and CL (Fig 3a). Traps baited with CL caught 83.8% of all specimens, while ZN and BL attracted 13.2% and 3% of fruit flies respectively. *Zeugodacus cucurbitae* was the most abundant species in CL traps followed by *D. telfaireae and D. punctatifrons* Karsch (Fig 3b). On the other hand, *D. durbanensis* was the most abundant species in ZN traps followed by *Z. cucurbitae* and D. *humeralis*. Specimens found in BL traps were dominated by *Z. cucurbitae* followed by *Dacus bivittatus* (Bigot) and *D. telfaireae* (Fig 3c).

**Fig 3a.**
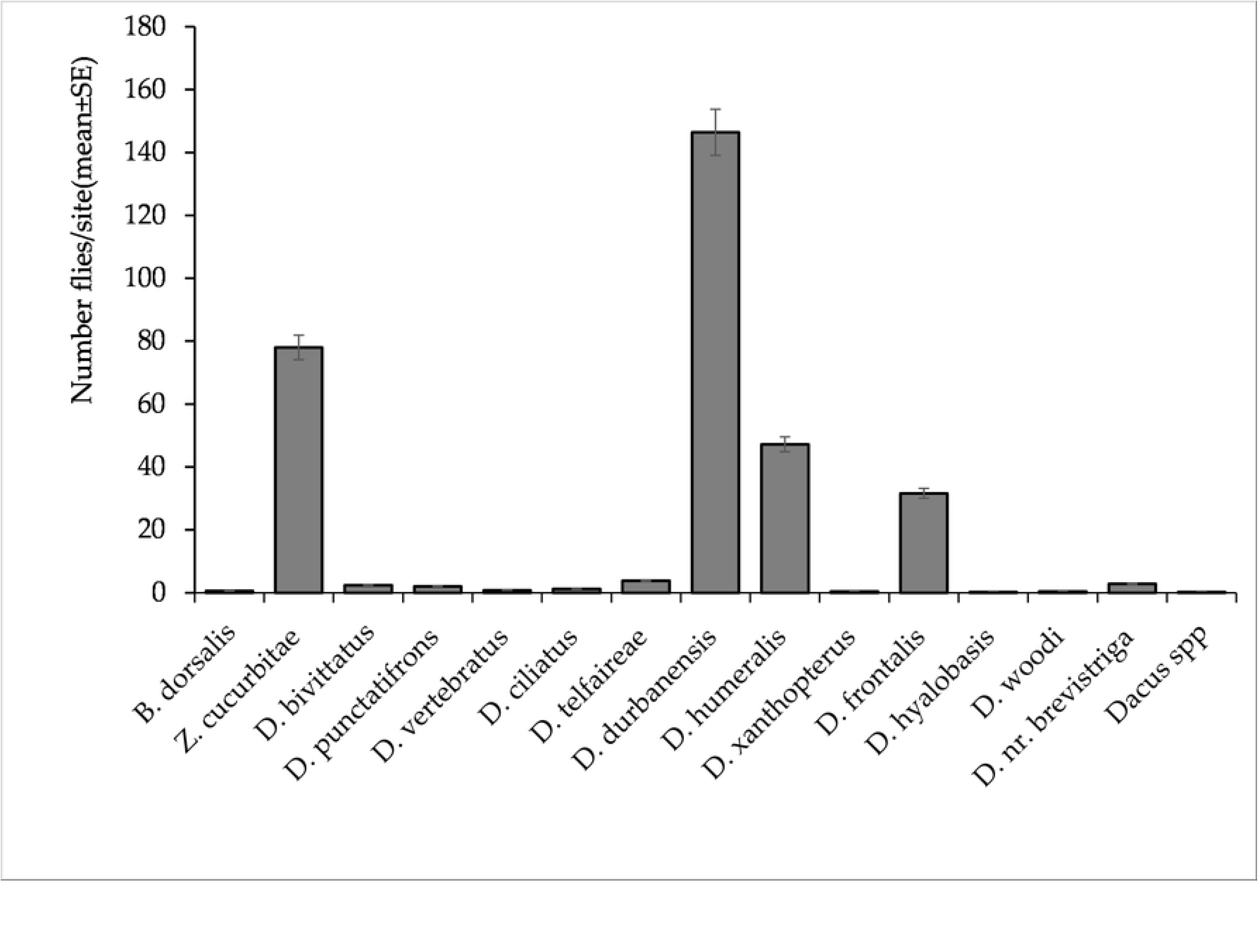
The abundance of cucurbit infester species trapped by Zingerone in the two agroecological zones of Morogoro, Tanzania from March to October 2020.

**Fig 3b.**
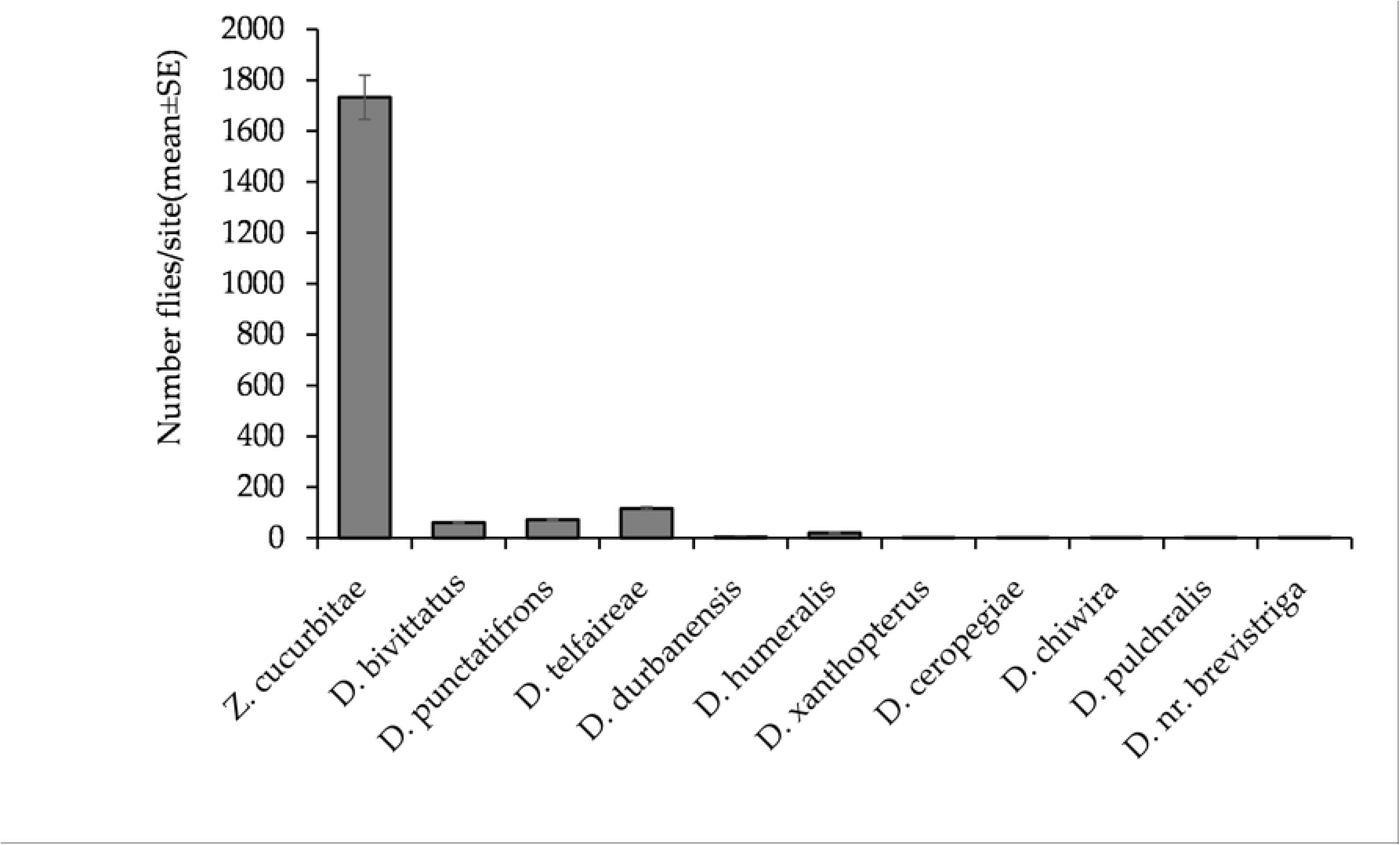
The abundance of cucurbit infester species trapped by Cue lure in the two agroecological zones of Morogoro, Tanzania from March to October 2020.

**Fig 3c.**
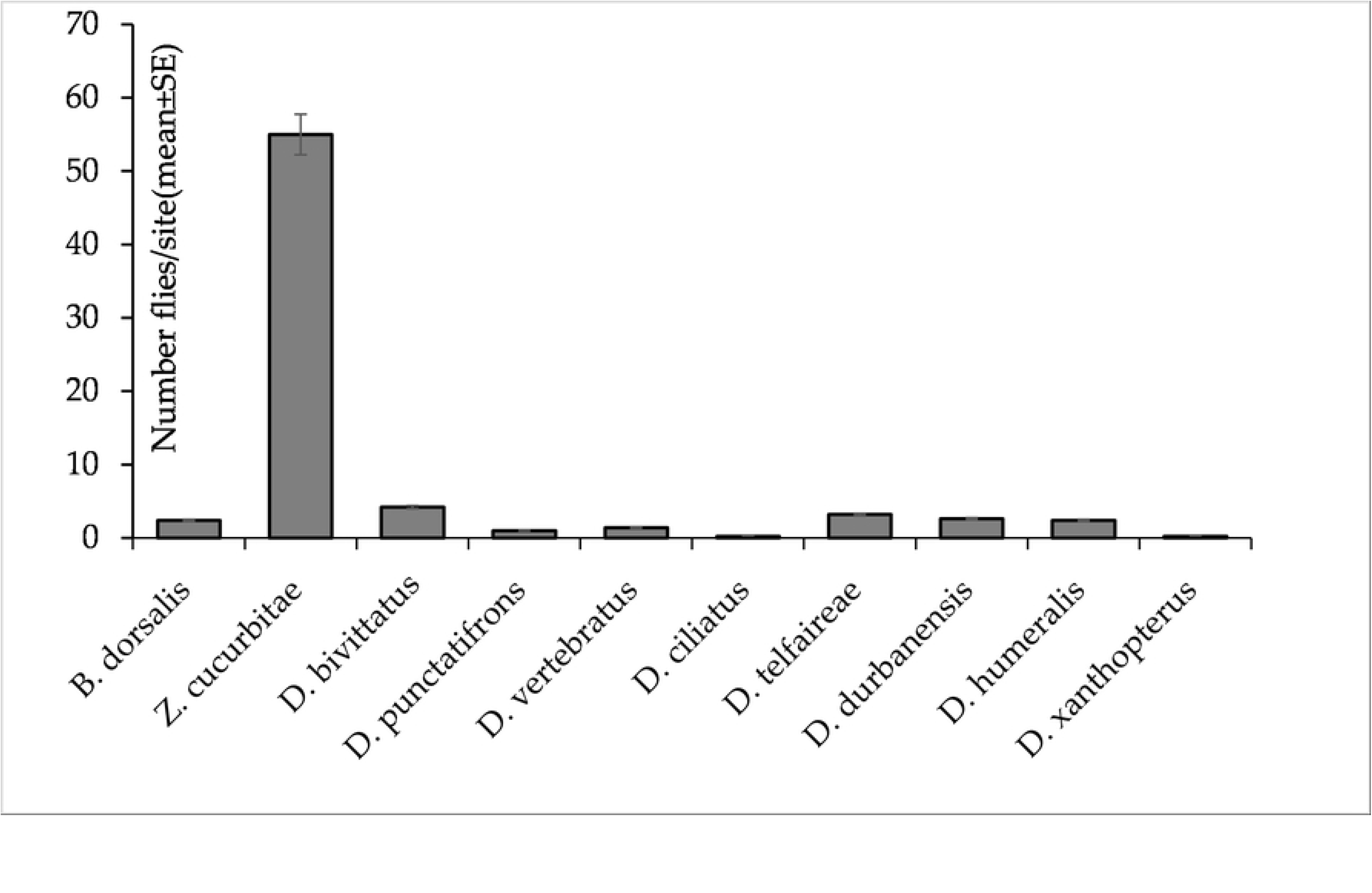
The abundance of cucurbit infester species trapped by Bio lure in the two agroecological zones of Morogoro, Tanzania from March to October 2020.

### 3.3. Within habitat diversity of fruit flies (Alpha diversity)

Results showed that fruit fly species richness, evenness, and diversity (Shannon’s and Inverse Simpson’s indices) from traps were higher in the mountainous than plateau zone (Table 4). These differences in richness, evenness, and diversity between the two agroecological zones were significant (Table 5, Figs 4a – d). We also found significant differences in richness, evenness, and diversity between the two cropping seasons. However, the effects of agroecological zone × season were not significant. Species richness, evenness and diversity were significantly higher during the June-August than September-November cropping season (Figs 4a – d).

**Table 4.** Mean alpha diversity indices values.

| Agroecological zone | Richness | Evenness | Shannon | Inverse Simpson |
| --- | --- | --- | --- | --- |
| **Mountainous** | 17±0.170 | 0.778±0.043 | 0.763±0.416 | 1.988±0.111 |
| **Plateau** | 11±0.126 | 0.654±0.041 | 0.521±0.396 | 1.444±0.112 |

**Table 5.** Analysis of variance for the effects of agroecological zones and Seasons on species richness, evenness, and diversity of fruit flies.

| Variables | Source of Variations | Statistics |  |
| --- | --- | --- | --- |
|  |  | *F value* | *P-value* |
| **Richness** | Agroecological zones | 8.313 | 0.004* |
|  | Season | 5.673 | 0.018* |
|  | Agroecological zone × season | 0.601 | 0.439ns |
| **Evenness** | Agroecological zone | 4.728 | 0.031* |
|  | Season | 9.364 | 0.003* |
|  | Agroecological zone × season | 0.301 | 0.584ns |
| Shannon | Agroecological zone | 9.169 | 0.003* |
|  | season | 7.644 | 0.006* |
| | Agroecological zone × season | 0.953 | 0.330$\mathbf{ns}$ |
| Inverse Simpson | Agroecological zones | 12.126 | 0.001* |
|  | Season | 4.558 | 0.034* |
| | Agroecological zone × Season | 1.460 | 0.229$\mathbf{ns}$ |

Note: values with \* are significant at $P < 0.05$ otherwise $\mathbf{ns}$ implies not significant.

**Fig 4a.**
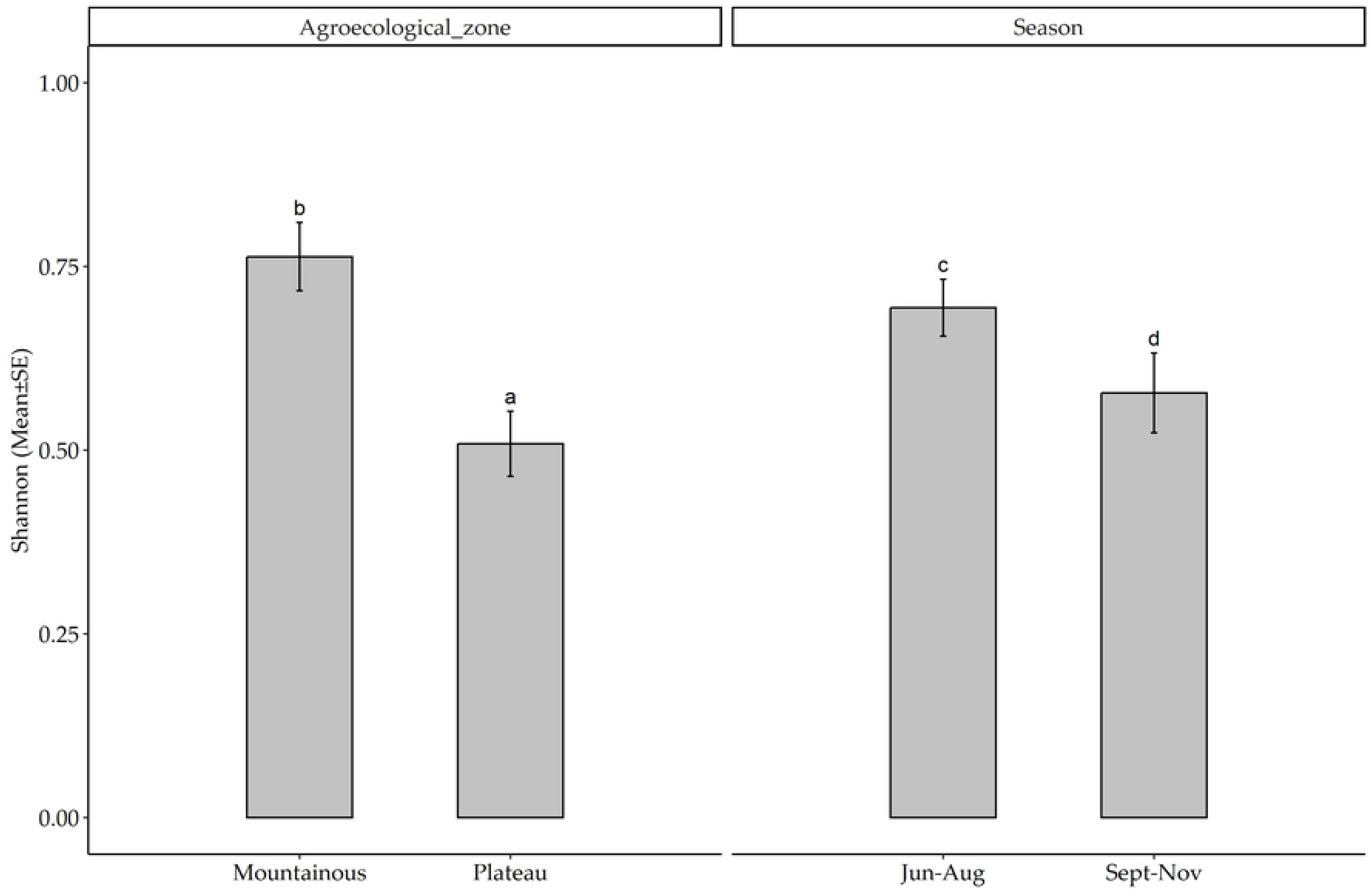
Shannon diversity of fruit fly species infesting cucurbit crops across the agroecological zones of Morogoro, Tanzania from June to November 2020. Bars with different letters denote significant differences, p < 0.05. SE stands for Standard Error.

**Fig 4b.**
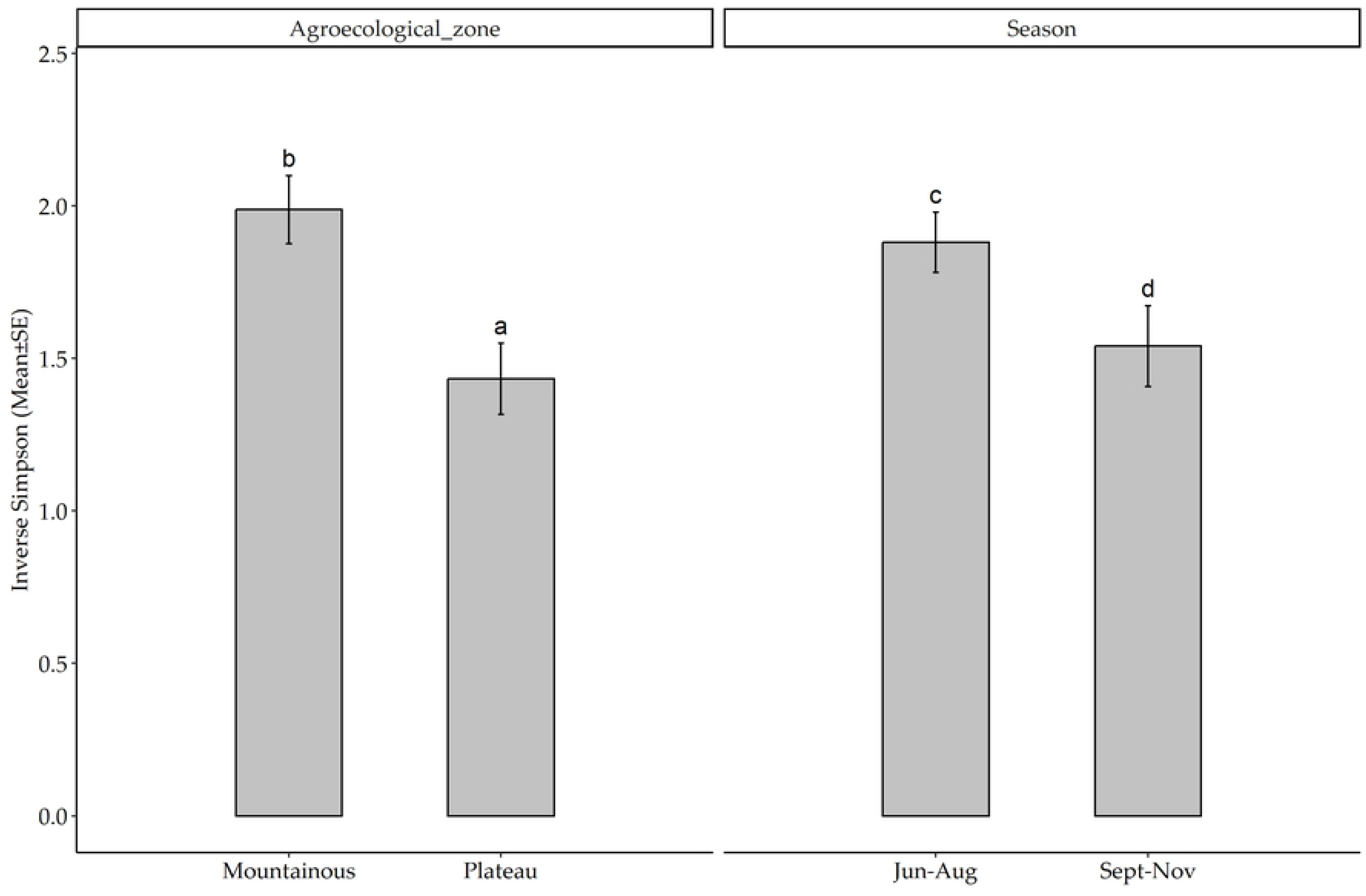
Inverse Simpson diversity of fruit fly species infesting cucurbit crops across the agroecological zones of Morogoro, Tanzania from June to November 2020. Bars with different letters denote significant differences, p < 0.05. SE stands for Standard Error.

**Fig 4c.**
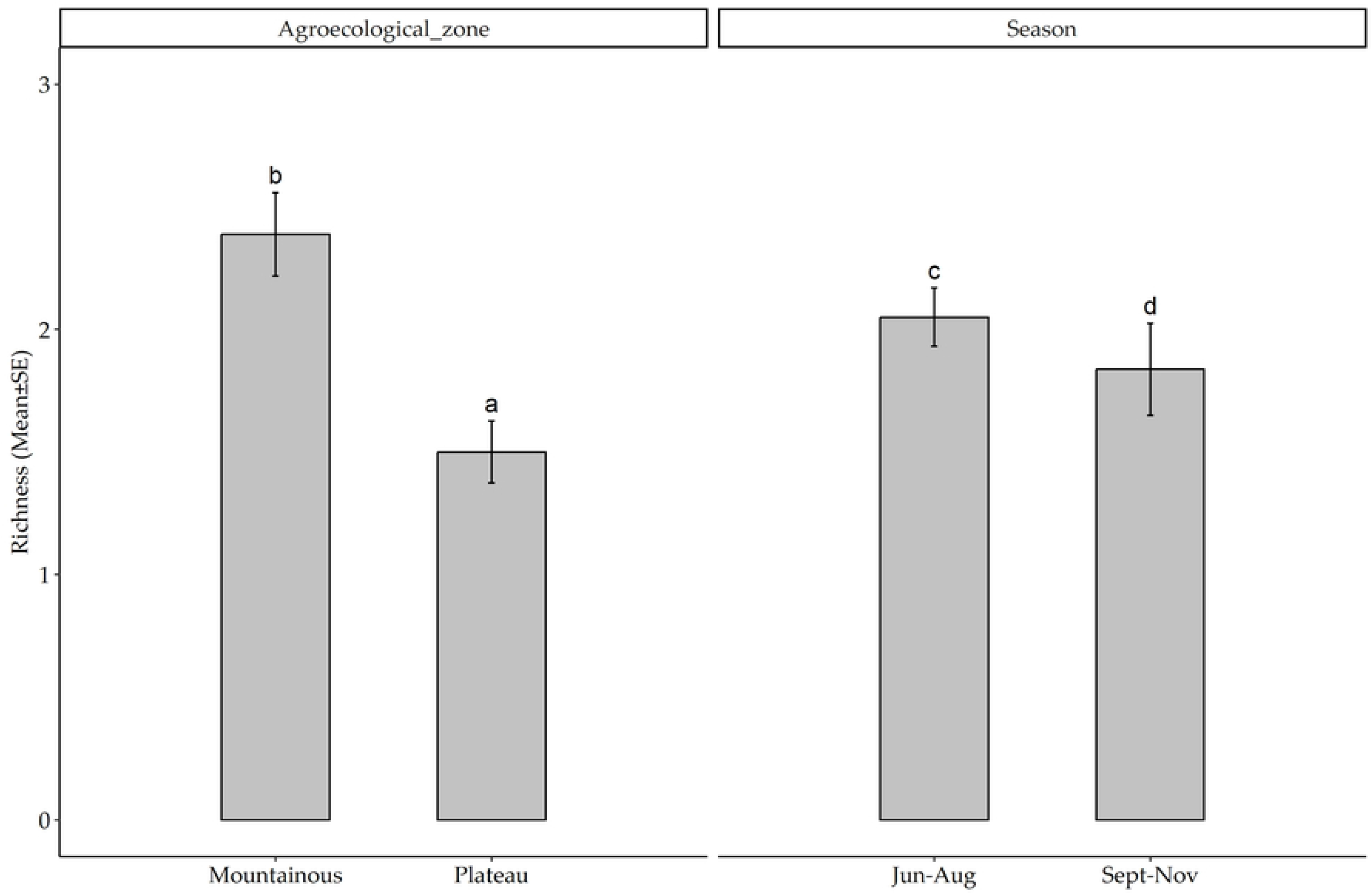
Fruit fly species richness infesting cucurbit crops across the agroecological zones of Morogoro, Tanzania from June to November 2020. Bars with different letters denote significant differences, p < 0.05. SE stands for Standard Error.

**Fig 4d:**
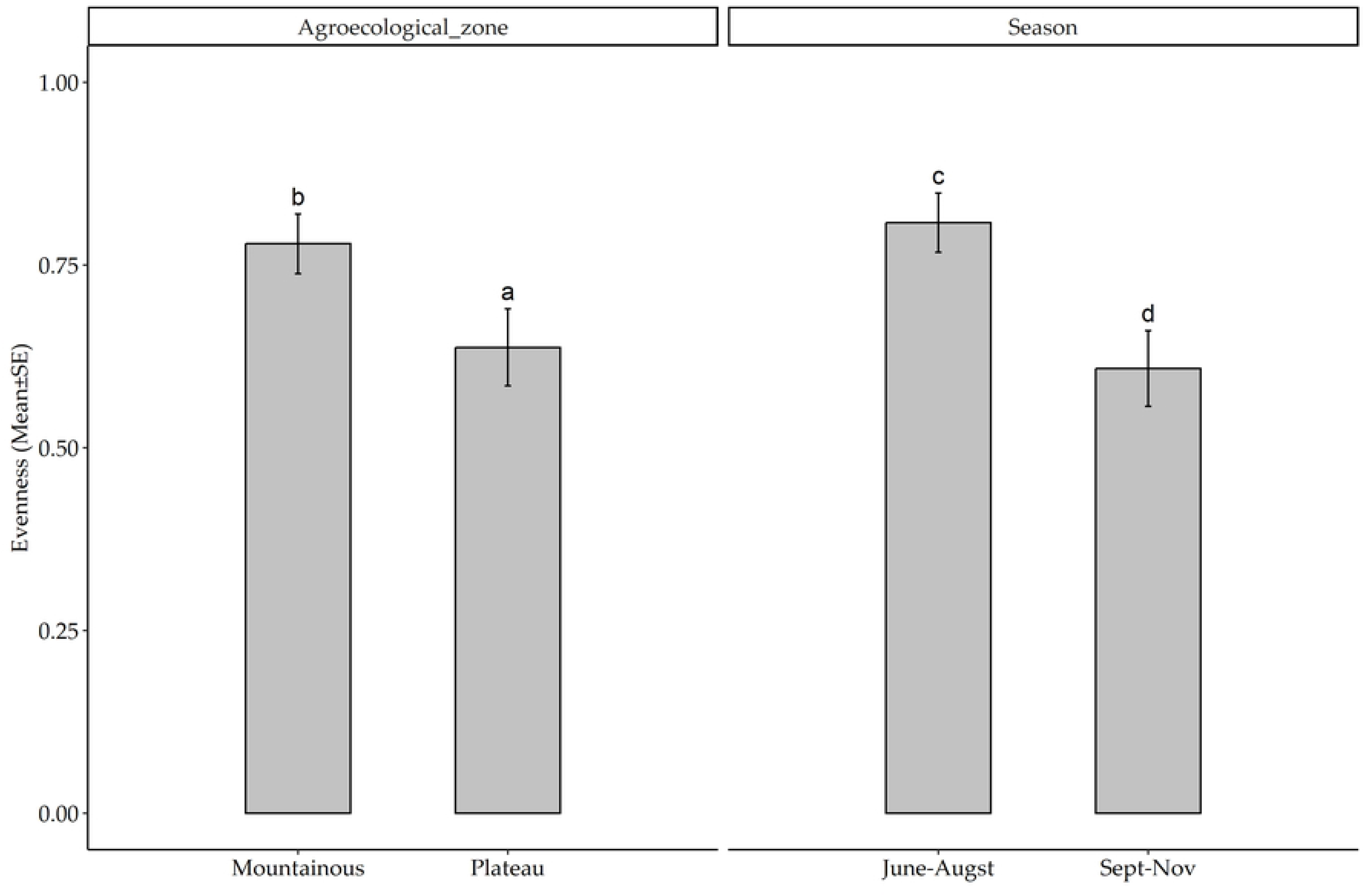
Fruit fly species evenness infesting cucurbit crops across the agroecological zones of Morogoro, Tanzania from June to November 2020. Bars with different letters denote significant differences, p < 0.05. SE stands for Standard Error.

### 3.4. Between habitats diversity of fruit flies (Beta diversity)

Both Sorensen’s and Jaccard’s indices had low values (Table 6), indicating a considerably high dissimilarity in terms of species composition between the plateau and the mountainous zone (Table 6).

**Table 6.** Beta diversity indices comparing mountainous and plateau zone.

| Index | Value |
| --- | --- |
| Sorensen (S) | 0.331 |
| Jaccard (J) | 0.201 |

### 3.5. Seasonal abundance of dominant fruit fly species

Out of ten fruit fly species caught by traps baited by Bio-lure, Cue-lure and Zingerone, *Z. cucurbitae, D. bivittatus* and *D. punctatifrons* were the first three most abundant species and major prominent pests of vegetable fruits known in the Morogoro region [53, 52]. Therefore, these species were used for further analysis.

Results from Figs 5a-c show that *Z. cucurbitae* dominates the Morogoro region with one to two distinct peaks throughout the two cucurbit cropping seasons. The other two species followed the same ‘ups and downs’ trends as *Z. cucurbitae* but with lower numbers with the exception of the *D. bivitattus* caught by traps baited with Cue-lure in the mountainous zone. *Z. cucurbitae* was recorded in higher numbers in traps baited with Bio lure and Zingerone in October and November (Figs 5a-b). For traps baited with Cue-lure, a higher number of *Z. cucurbitae* and *D. bivittatus* per week were recorded during the end of July and October as well as during early August and November (Fig 5c). The other two remaining species maintained similar fluctuation trends but in relatively low numbers (Figs 5a-c).

**Fig 5a.**
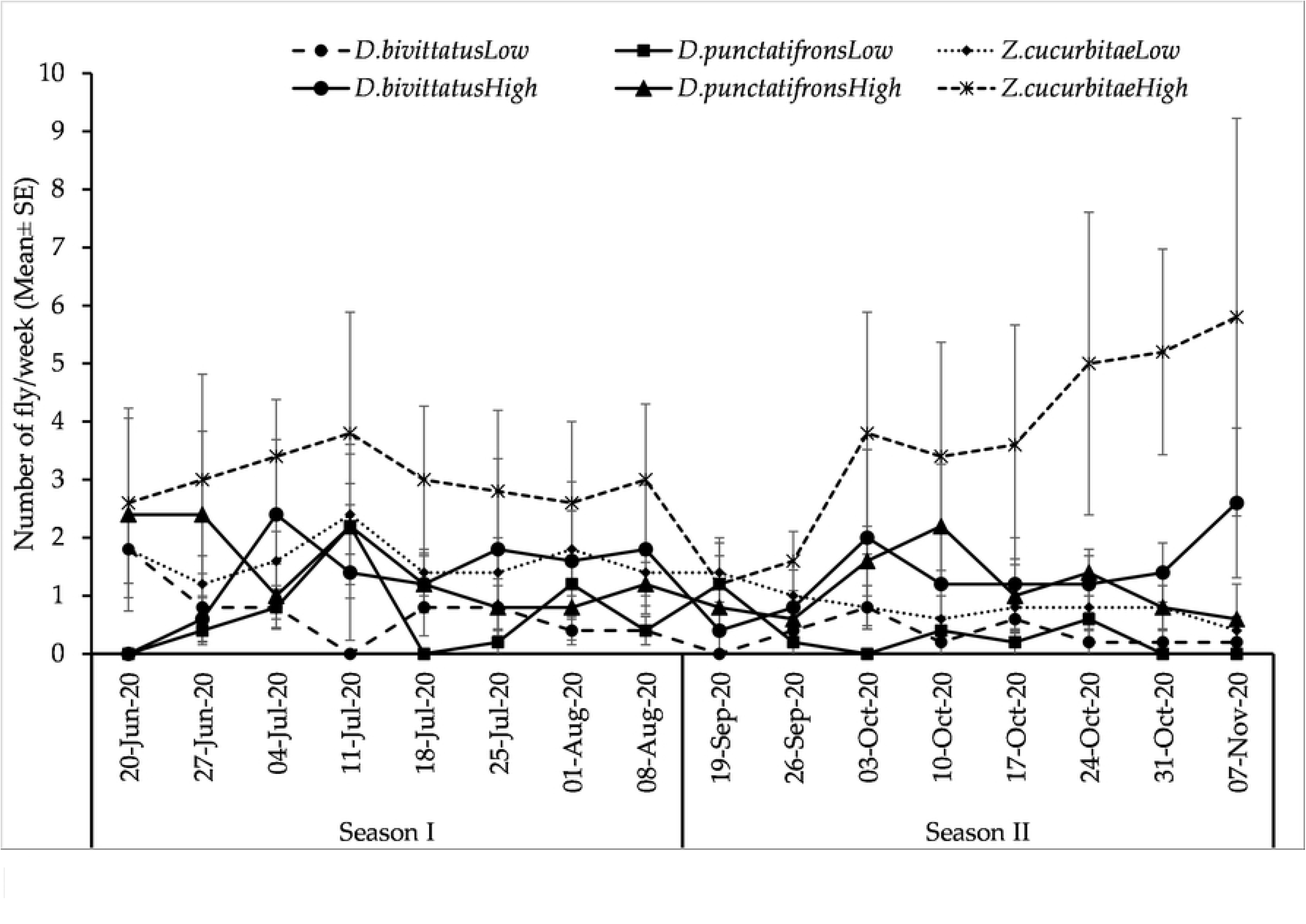
Seasonal fluctuation of most abundant fly species infesting cucurbit crops in the Morogoro region trapped using Biolure from June to November 2020. SE stands for Standard Error.

**Fig 5b.**
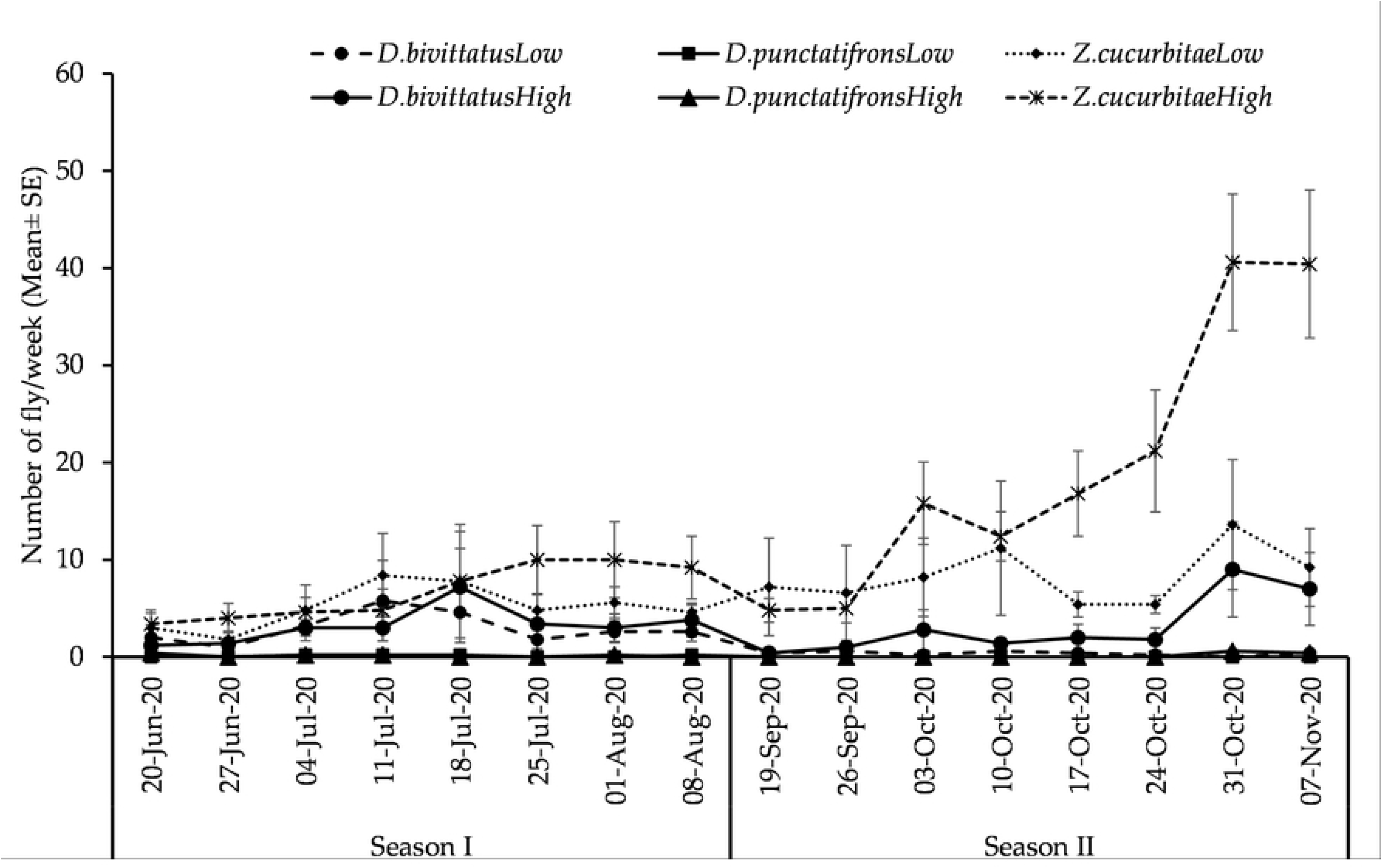
Seasonal fluctuation of most abundant fly species infesting cucurbit crops in the Morogoro region trapped using Zingerone from June to November 2020. SE stands for Standard Error.

**Fig 5c.**
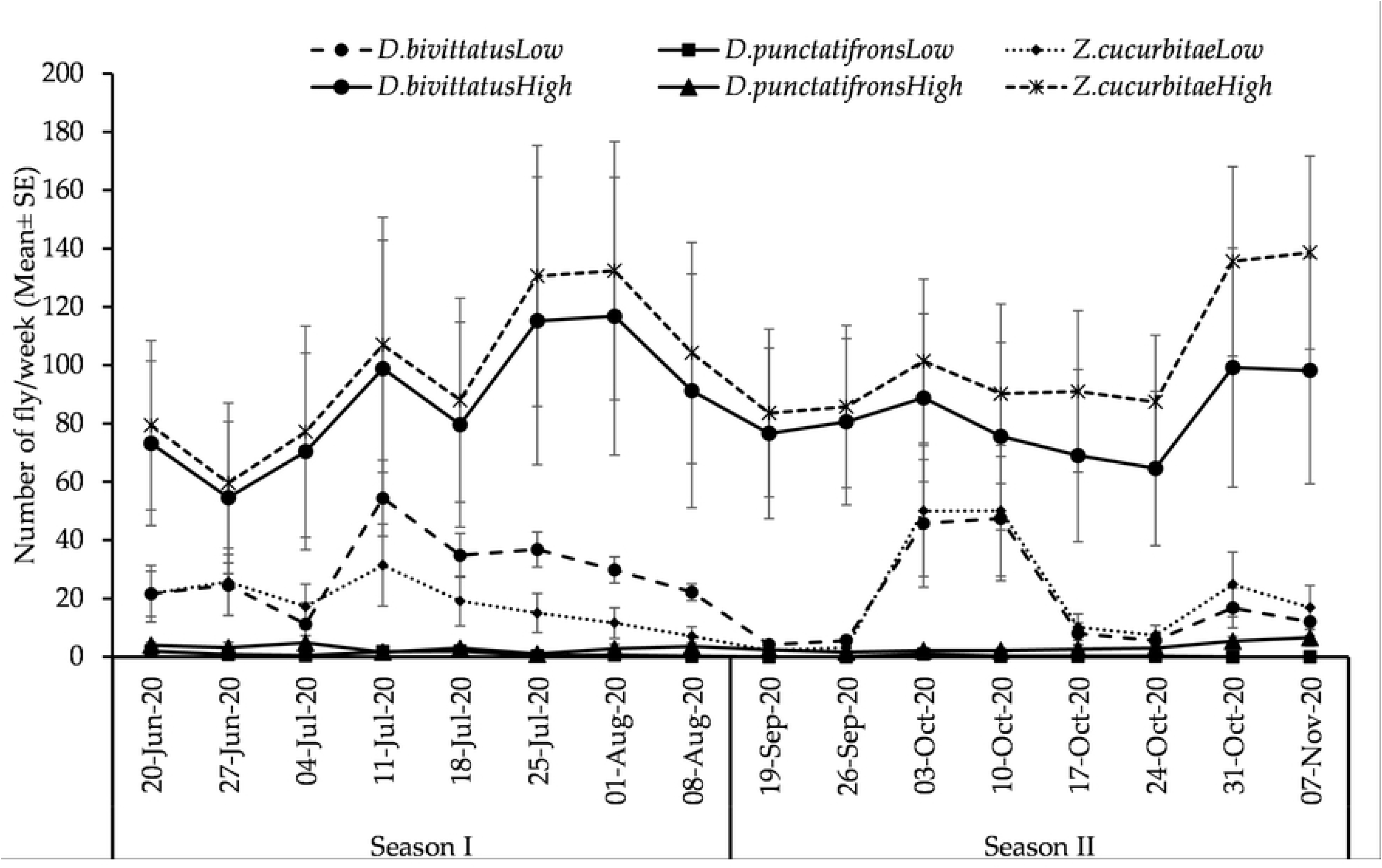
Seasonal fluctuation of most abundant fly species infesting cucurbit crops in the Morogoro region trapped using Cue-lure from June to November 2020. SE stands for Standard Error.

### 3.6. Effect of agroecological zones and seasons on the abundance of most abundant fruit fly species

The ANOVA results on the effect of agroecological zones fly species and cropping seasons on the abundance of fruit fly species are presented in Table 7. Results showed that the abundance of a species varied significantly (*p* < 0.05) in the agroecological zone as well as with cropping season. Further examination of results, showed significant variations within each agroecological zone (Fig 6) and season (Fig 7). The abundance of *Z. cucurbitae* was significantly higher than the other species at both altitudes and cropping seasons (*Post hoc* test = Student-Newman-Keuls). The effects of the interaction between fly species, season and altitude were not significant (p > 0.05).

**Fig 6.**
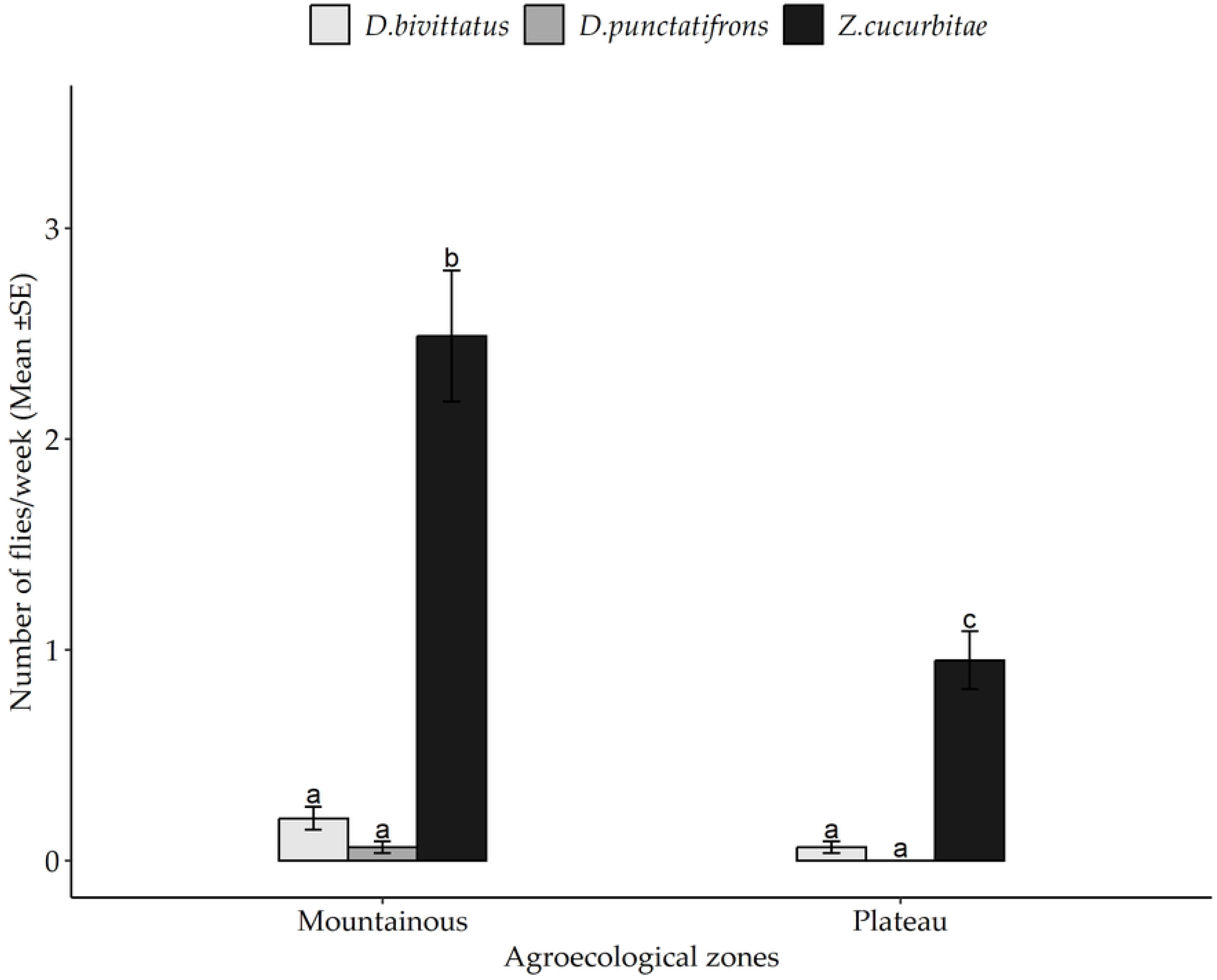
The abundance of dominant fruit fly species across the two agroecological zones. Bars with different letters denote significant differences, p < 0.05. SE stands for Standard Error.

**Fig 7.**
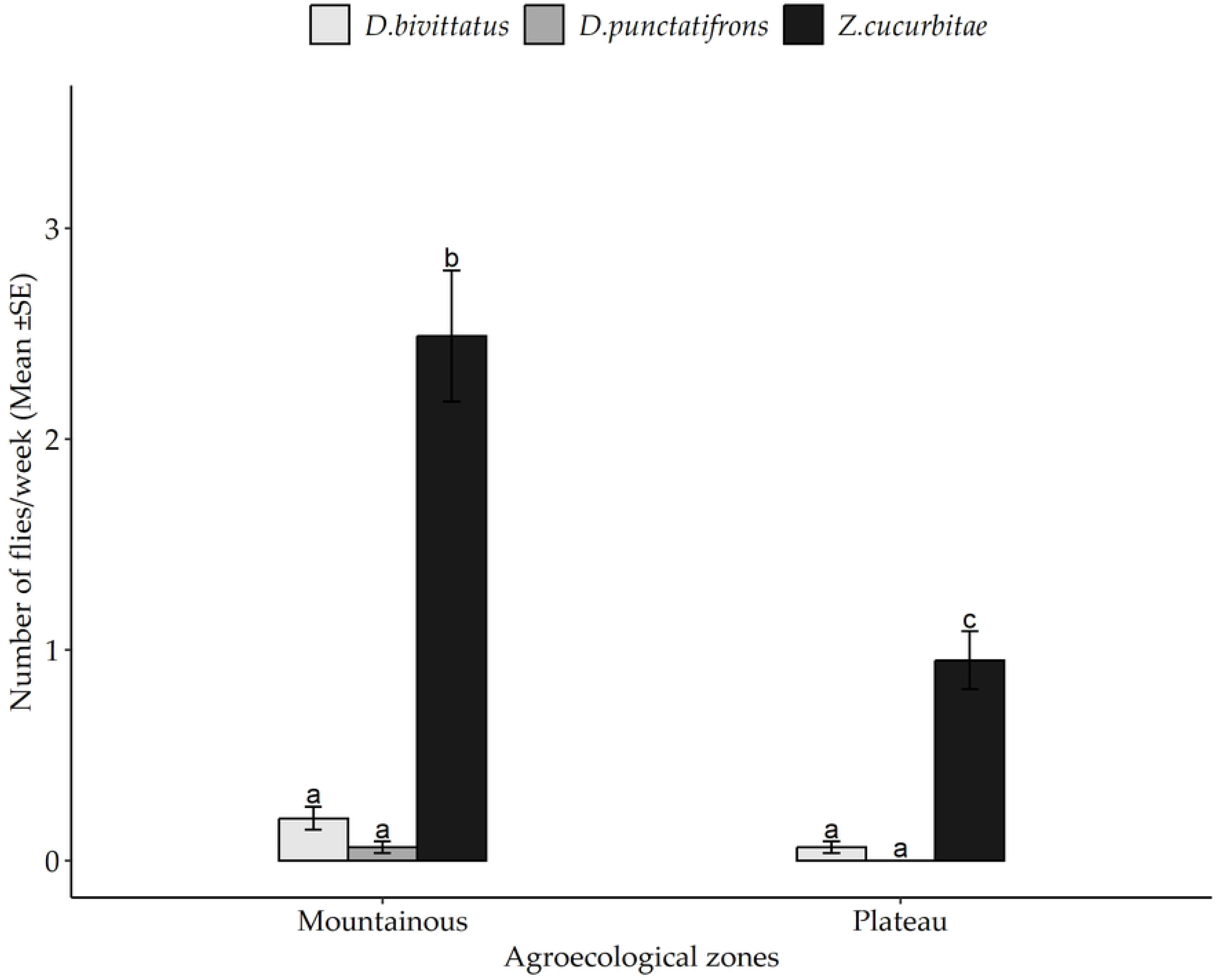
The abundance of dominant fruit fly species between two cucurbit cropping seasons. Bars with different letters denote significant differences, p < 0.05. SE stands for Standard Error.

**Table 7.**
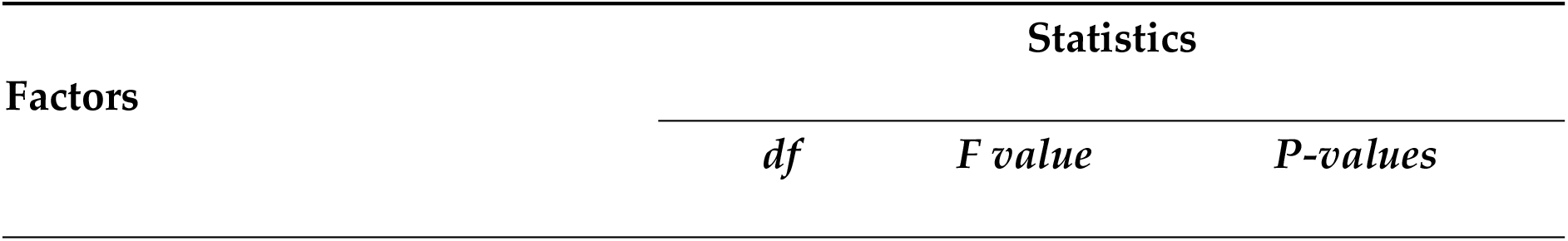
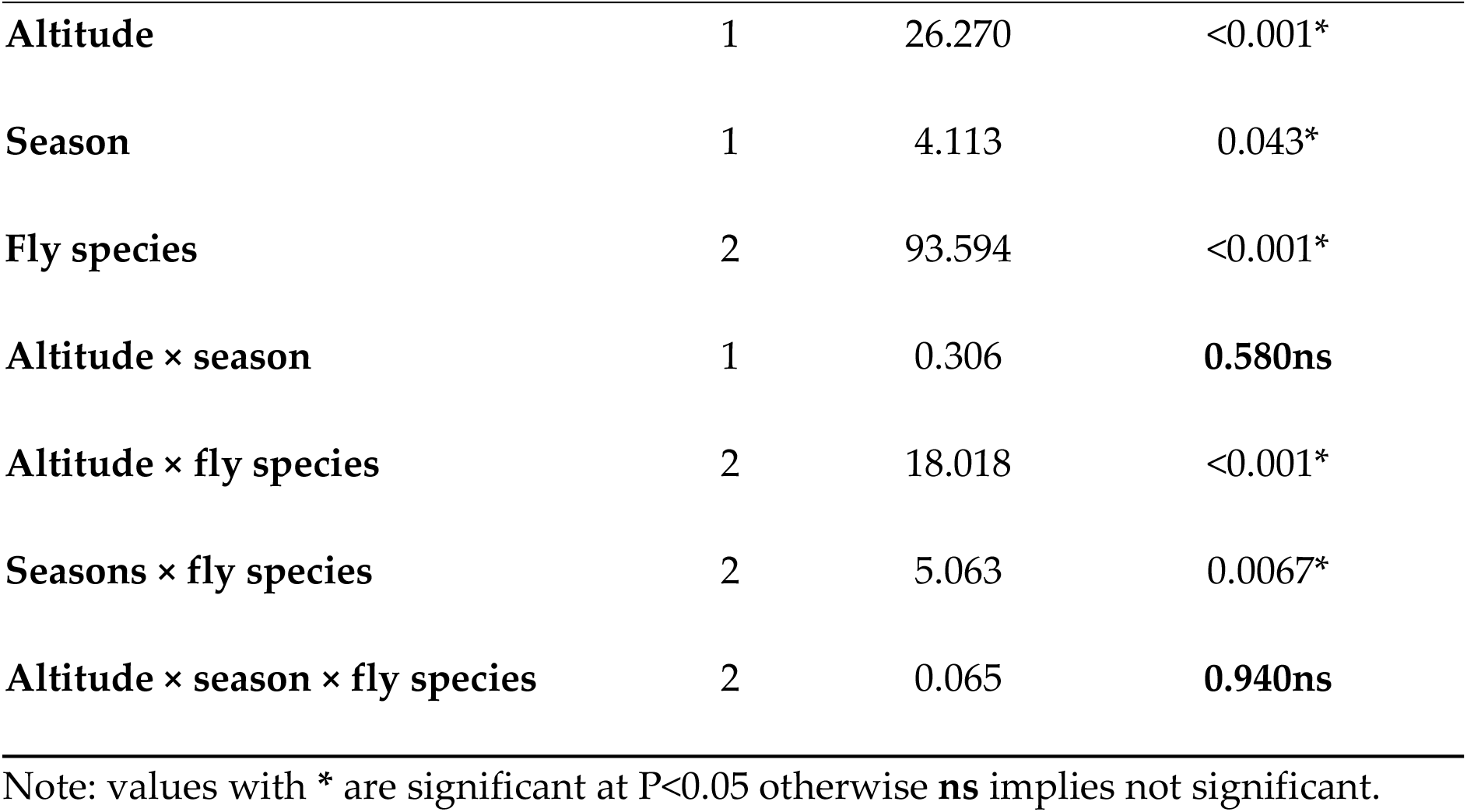
Analysis of variance for the effects of agroecological zones, seasons, and Fly species on the weekly abundance of fruit flies.

| Factors | Statistics |  |  |
| --- | --- | --- | --- |
|  | *df* | *F value* | *P-values* |
| **Altitude** | 1 | 26.270 | <0.001* |
| **Season** | 1 | 4.113 | 0.043* |
| **Fly species** | 2 | 93.594 | <0.001* |
| **Altitude × season** | 1 | 0.306 | **0.580ns** |
| **Altitude × fly species** | 2 | 18.018 | <0.001* |
| **Seasons × fly species** | 2 | 5.063 | 0.0067* |
| **Altitude × season × fly species** | 2 | 0.065 | **0.940ns** |

Note: values with \* are significant at $P < 0.05$ otherwise **ns** implies not significant.

## 4. Discussion

### 4.1. Sampling effort

Sampling effort is highly important when designing a biodiversity study to generate an inventory list of species of a locality for management purposes [33, 43]. It provides information on the minimum number of sites to be sampled per zone and the number of samples required to ensure satisfactory representativeness of a true community under investigation [4]. Results of the three nonparametric estimators of species richness revealed that the sampling effort was adequate to recover more than 85 % of fruit fly species in all agro-ecological zones. Based on SACs results, about 9700 and 3200 individuals were recovered in the mountainous and plateau zone respectively from 360 samples collected per zone. Both samples were reasonable and acceptable for the study. According to [1], conventionally the minimum sample size accepted from optimal sampling and efficient comparisons of ecological communities is 20 samples. Thus, the sampling effort determined in the present study was sufficient to obtain a satisfactory representation of samples of fruit fly communities.

### 4.2. Diversity of fruit flies

The results revealed significant differences in fruit fly species richness, evenness, and diversity between the two agroecological zones of the Morogoro region and between the two-cropping season. The mountainous zone was numerically the most diverse and species-rich compared to the plateau zone. The mountainous zone harboured a variety of known cucurbit infesters (*Z. cucurbitae*, *D.bivittatus, D.punctatifrons, D.ciliatus* and *D.telfaireae*) and other non-cucurbit feeders (*D. durbanensis, D. humeralis, D. xanthopterus, D. frontalis, D. hyalobasis, D. hyalobasis, D. woodi, D. ceropegiae, D. chiwira, D. pulchralis* and *D. nr. brevistriga*) than the plateau.

The difference in the diversity of tephritid species including non-cucurbit feeders between the two agroecological zone can be attributed to the availability of wild host plants in the adjacent areas. The experimental plots in the mountainous zone were located proximal to the natural forest of Uluguru mountains, where wild hosts are abundant compared to the plateau zone. The observed variation in fruit fly diversity could also be related to differences in predominant farming practices between the agroecological zones. The mountainous landscape is predominated by agroecological farming practices [11, 40, 51]. These could favour higher diversity compared to the plateau zone which is largely in a landscape that is predominated by conventional agriculture [56, 61]. Seasonal shifts in the availability of fruit hosts could result in higher fruit fly abundance or an increase in rare species depending on the season [53]. In this case, host availability is the driving factor for seasonal fluctuations of fruit fly population density [21, 22, 47]. In the tropics, fruit fly seasonality is often linked to hosts that may serve as a bridge allowing populations of fruit flies to persist at times of the year when the main hosts are not available [21, 41, 47] resulting in host shifts depending on the season [53].

The frequent encounter of *Z. cucurbitae, D. punctatifrons* and *D. bivittatus* across the ecological zones as dominant agricultural pests have been also reported in different parts of Africa [2, 15, 28, 42, 49]. The other species from the genus *Dacus* were also found but in low numbers, often represented by very few specimens per field.

### 4.2. Abundance of cucurbit infesters

The spatial and temporal abundance of selected cucurbit infesters in the Morogoro region was significantly affected by the effects of season, agroecological zone and fruit fly species. The results also showed that the abundance of *Z. cucurbitae* was higher than *D. bivittatus* and *D.punctatifrons* both at the plateau and mountainous zones. There are mixed reports on the abundance of *Z. cucurbitae* across altitudes and in presence of other fruit fly species. In the Reunion islands, the relative abundance of *Z. cucurbitae* was lowest in high-altitude sites (above 1000 m), where *D. demmerezi* was the most prevalent species [62]. In this case, *Z. cucurbitae* was found at low and medium altitudes competing with *D. ciliatus.* A later study by [14] on Reunion islands reported that *Z. cucurbitae* was the least abundant species compared to *D. ciliatus* and *D. demmerezi*. According to [48] the occurrence of *Z. cucurbitae* in the Morogoro Region was limited to medium altitude due to its preference for warmer conditions. However, the previous studies in Morogoro were based on trapping in fruit orchards and random sampling of both wild and cultivated hosts. In the present study, commercial hosts were purposely established and maintained under irrigation for extended periods of the year.

Our results also showed that *Z. cucurbitae* was the dominant species regardless of the cropping season. The abundance was higher during the June – August period than the September – November period. A previous study by [41, 48] in Morogoro reported that the peak populations of *Z. cucurbitae* were also recorded during the dry period when temperatures and relative humidity were low, but when hosts were widely available. On the contrary, [32] found that *Z. cucurbitae* was more active during warm and rainy months as compared to dry and winter months. It can be inferred that multiple factors, apart from the weather, modulate the population of *Z. cucurbitae* [31].

## 5. Conclusion

The study successfully characterized the fruit fly communities in the Morogoro region across the plateau and mountainous zone. The species diversity was higher in the mountainous zone than in the plateau zone. Furthermore, *Z. cucurbitae* was the most abundant species in both zones. Therefore, this study highlights the need for considering agroecological zones, availability of host crops and proximity of natural vegetation during the designing of any sustainable control strategy against fruit flies. Management programs against cucurbit infesters should be more intensive and centred on *Z. cucurbitae, D. punctatifrons* and *D. bivittatus* regardless of the agroecological zones. Both short and dry seasons should also be considered since the results of this study showed a higher abundance of *Z. cucurbitae* during the short rain and dry seasons.

## Acknowledgements

Thanks are due to Cessila Kijalo, Evance Matowo, Frida Mangu, Christian Barnabas and Tabitha Fussi (Sokoine University of Agriculture) for their assistance in the field and Laboratory work, to I.M. White (Natural History Museum, London) for making available his forthcoming revision on dacines of the Afrotropical region. We also extend our thanks to Wouter Hendrycks and Lore Esselens for assisting analysis of data.

